# Comparison of colorimetric, fluorometric, and liquid chromatography-mass spectrometry assays for acetyl-coenzyme A

**DOI:** 10.1101/2023.06.01.543311

**Authors:** Daniel S. Kantner, Emily Megill, Anna Bostwick, Vicky Yang, Carmen Bekeova, Alexandria Van Scoyk, Erin Seifert, Michael W. Deininger, Nathaniel W. Snyder

**Affiliations:** Lewis Katz School of Medicine at Temple University, Department of Cardiovascular Sciences, Center for Metabolic Disease Research, Philadelphia, PA 19140, USA; MitoCare Center, Department of Pathology, Anatomy, and Cell Biology, Thomas Jefferson University, Philadelphia, PA 19107, USA; Huntsman Cancer Institute, University of Utah, Salt Lake City, UT 84112, USA; Versiti Blood Research Institute and Medical College of Wisconsin, Milwaukee, WI 53226, USA

**Keywords:** Mass spectrometry, acetyl-CoA, metabolite, method comparison

## Abstract

Acetyl-Coenzyme A is a central metabolite in catabolic and anabolic pathways as well as the acyl donor for acetylation reactions. Multiple quantitative measurement techniques for acetyl-CoA have been reported, including commercially available kits. Comparisons between techniques for acetyl-CoA measurement have not been reported. This lack of comparability between assays makes context-specific assay selection and interpretation of results reporting changes in acetyl-CoA metabolism difficult. We compared commercially available colorimetric ELISA and fluorometric enzymatic-based kits to liquid chromatography-mass spectrometry-based assays using tandem mass spectrometry (LC-MS/MS) and high-resolution mass spectrometry (LC-HRMS). The colorimetric ELISA kit did not produce interpretable results even with commercially available pure standards. The fluorometric enzymatic kit produced comparable results to the LC-MS-based assays depending on matrix and extraction. LC-MS/MS and LC-HRMS assays produced well-aligned results, especially when incorporating stable isotope-labeled internal standards. In addition, we demonstrated the multiplexing capability of the LC-HRMS assay by measuring a suite of short-chain acyl-CoAs in a variety of acute myeloid leukemia cell lines and patient cells.

## Introduction

Acetyl-coenzyme A (acetyl-CoA) functions as a central metabolite and the acyl-donor for post-translational acetylation of proteins and other biological molecules [1]. Consequently, acetyl-CoA is an important biological molecule to consider when evaluating the metabolic and energetic status of cells. Acetyl-CoA metabolism constitutes a drug target actively under investigation for multiple diseases including one approved drug (bempedoic acid) for cholesterol and lipid reduction with demonstrated benefit [2]. Therefore, sensitive and specific techniques for quantitating acetyl-CoA in various biological samples are of interest to researchers with diverse biological interests. Liquid chromatography-mass spectrometry-based assays for acetyl-CoA (and other CoA derivatives) have been developed, validated, and applied in different contexts, with liquid chromatography-tandem mass spectrometry (LC-MS/MS), liquid chromatography-high resolution mass spectrometry (LC-HRMS), direct injection HRMS, and capillary electrophoresis-MS methods reported [3-13].

A limited number of commercial acetyl-CoA assay kits are available, based on reagent-specific detection by colorimetric or fluorometric readouts. LC-MS varies by the specifics of instrumentation and is often found in specialized laboratories, but UV/vis plate readers with appropriate filters are more widely distributed instrumentation. Thus, commercial acetyl-CoA assay kits coupled with colorimetric and fluorescent detection provide some laboratories with a more accessible method of acetyl-CoA quantitation. However, to date, no comparisons between LC-HRMS, LC-MS/MS methods, and commercial kits have been documented.

We conducted a comparison of a colorimetric acetyl-CoA ELISA, an enzymatic fluorometric acetyl-CoA assay, and LC-MS/MS and LC-HRMS assays of water-soluble short-chain acyl-CoAs. While the colorimetric ELISA kit did not produce interpretable results even with commercially available pure standards, the fluorometric enzymatic kit produced comparable results to the LC-MS-based assays depending on the matrix and extraction solution utilized. In addition, we demonstrated that both the fluorometric and LC-MS-based assays can be scaled to 96-well format with liquid and solid phase extraction and concentration procedures. LC-MS/MS and LC-HRMS assays produced comparable measurements, and both assays benefitted from the use of stable isotope-labeled internal standardization. Also, the LC-HRMS assay was determined to be suitable for the multiplexed analysis of a variety of water-soluble short-chain acyl-CoAs.

## Materials and Methods

### Chemicals and Reagents

Acetyl-CoA lithium salt (used as the LC-MS calibration standard) and 5-sulfosalicylic acid (SSA) were purchased from Sigma-Aldrich (P/N: A2181 and P/N: S2130, respectively). Optima® LC/MS grade acetonitrile (ACN), formic acid, methanol, and water were purchased from Fisher Scientific. High-glucose Dulbecco’s Modified Eagle Medium (DMEM) was purchased from Gibco. Oasis® HLB solid-phase extraction (SPE) columns and 96-well elution plates (30 mg of sorbent) were purchased from Waters (P/N: 186003908 and P/N: WAT058951, respectively). The acetyl-CoA ELISA kit was purchased from Elabscience (P/N: E-EL-0125). The PicoProbe^™^ acetyl-CoA fluorometric assay kit was purchased from BioVision/abcam (P/N: K317 / ab87546). The perchloric acid (PCA) deproteinizing sample preparation kit was purchased from BioVision/abcam (P/N: K808 / ab284939). The short-chain acyl-CoA internal standard for the LC-MS assays was generated in yeast as previously described [14].

### Equipment

A Fisherbrand^™^ Sonic Dismembrator (Model 120) equipped with a single-tip Qsonica CL-18 or a 602-A 8-tip sonicator probe was used to perform sonication. The ELISA samples were analyzed with a BioTek Synergy LX plate reader set to 450 nm. The PicoProbe™ assay samples were analyzed with a BioTek Synergy LX plate reader installed with a red filter cube (Excitation/Emission: 530/590 nm; P/N: 1505004).

### Cell Culture and Tissue Samples

All cells were cultured at 37°C with 5% CO_2_. HepG2, HAP1, and Panc-1 SLC25A20 cells were cultured in DMEM supplemented with 10% fetal bovine serum (FBS) and 1% penicillin/streptomycin. For the method comparison experiments, cells were allowed to reach not less than approximately 80% confluence prior to harvesting for metabolite extraction.

Acute myeloid leukemia (AML) cell lines (CMK, GFD-8, HEL, K562, Kasumi-1, KBM3, KBM5, KCL22, KG1a, KU812, KY0-1, LAMA-84, M07e, MOLM-13, MOLM-14, Monomac-1, NOMO-1, OCI-AML2, OCI-AML3, OCI-AML5, SET2, SKM-1, SKNO-1, TF-1, THP-1) were grown in RPMI supplemented with 10% FBS (Sigma), 1% Glutamax (ThermoFisher), and 100 U/mL penicillin/streptomycin (ThermoFisher). MV-411 and UT-7 were grown in IMDM or MDM supplemented with 10% FBS, 1% Glutamax, and 100 U/mL penicillin/streptomycin. GFD-8, M07e, OCI-AML5, SKNO-1, TF-1, and UT-7 were supplemented with 5-50 ng/mL of GM-CSF. Primary AML cells were cultured in RPMI supplemented with 10% FBS, 1% Glutamax, 10 ng/mL GM-CSF, and 100 U/mL penicillin/streptomycin. Cell lines were authenticated using the GenePrint 24 kit (Promega) and DSMZ Online STR Analysis database at the DNA Sequencing Core at the University of Utah. All cell lines were confirmed for mycoplasma negativity using the MycoAlert Mycoplasma Detection Kit (Lonza). 24 hours before harvesting, cells were washed with Dulbecco’s phosphate-buffered saline (DPBS) and placed in fresh media. For harvesting, cells were washed three times with cold DPBS, spun down at 300 x g for 5 minutes at 4°C, and stored at -80°C until processing for LC-MS analysis.

Frozen mouse heart and skeletal muscle tissue samples were obtained from 11-to 13-week-old male and female C57BL/6 J mice. Tissues were always harvested in the morning, snap frozen, and stored at -80°C until analysis. Mice were maintained on a 12–12 h light–dark cycle (lights on: 7:00 to 19:00) and ad libitum fed a standard diet (LabDiet 5001, Purina). Animal studies were approved by the Jefferson University IACUC protocol #01307.

### Cell Volume and Size Measurements

For AML cell lines, cells were pelleted at 250 x g for 5 minutes at 4°C. Cells were resuspended in 2 mL of StemPro Accutase Cell Dissociation Reagent (ThermoFisher) at room temperature for 30 minutes. Samples were diluted in DPBS and run on a Moxi Z Mini Automated Cell Counter [15]. Cell size of primary patient samples was determined using the Countess II FL Automated Cell Counter.

### Method Comparison of LC-HRMS, LC-MS/MS, and ELISA Assays

HepG2 cells were plated in 10-cm dishes at three different cell densities (0.1, 1, or 10 million cells per dish) and incubated overnight. After incubation, the dishes were placed on a slope on ice, and the medium was aspirated from each plate. 1 mL of ice-cold 10% (w/v) trichloroacetic acid (TCA) in water was added to each plate, and cells were scraped into microfuge tubes. The cell suspension was sonicated (5 x 0.5-second pulses at 50% intensity) to lyse cells, and the lysate was centrifuged at 17000 x g for 10 minutes at 4°C. The clarified extract was split between the ELISA and LC-MS methods (140 μL and 600 μL, respectively). The ELISA aliquot was neutralized (∼ pH 8) using potassium hydroxide (2 M and 10 M) and 1 M Tris-HCl (pH 8), and the ELISA was performed per manufacturer’s recommendations using 100 μL of the neutralized extract. A standard series was prepared using commercial acetyl-CoA material (Sigma-Aldrich) and analyzed in parallel with a standard series prepared using the standard provided by the manufacturer.

For the LC-MS aliquot, 50 μL of acyl-CoA internal standard was added prior to applying to Oasis HLB SPE columns for sample cleanup. The SPE columns were preconditioned with 1 mL of methanol, followed by equilibration with 1 mL of Optima water. The sample aliquots were loaded onto the column and then washed with 1 mL of Optima water. Acetyl-CoA was eluted with 1 mL of 25 mM ammonium acetate in methanol. Samples were evaporated to dryness under nitrogen (N_2_) and resuspended in 50 μL of 5% (w/v) SSA in water. 5 μL of each sample was injected for LC-HRMS or LC-MS/MS analysis.

For mouse heart and skeletal muscle tissue samples, 1 mL of ice-cold 10% TCA was added to approximately 10 mg (mid-level) or 30 mg (high-level) of tissue. Tissue samples were sonicated and centrifuged as described above. A 100-μL aliquot of the supernatant from the 10-mg tissue samples was diluted 1:10 in ice-cold 10% TCA to generate low-level samples equivalent to approximately 1 mg of tissue. Aliquoting, neutralization, and further processing of tissue samples was performed as described for the cell samples.

### Method Comparison of LC-HRMS, LC-MS/MS, and Fluorometric PicoProbe™ Assays

#### Whole-cell and Tissue Extraction

HepG2 cells were pelleted, resuspended in 1 mL of medium, and counted via a hemocytometer or a Beckman Coulter counter. Cells were pelleted and resuspended in 500 μL of either 80:20 methanol:water (-80°C), ice-cold MS buffer, or ice-cold 10% TCA (see detailed extraction protocols). For mouse heart tissue samples, 500 μL of extraction solution was added to the tissue (between 45 and 80 mg), and the tissue was sonicated to homogeneity via multiple rounds of 5 x 0.5 second pulses. Homogenized tissue samples were processed identically to the cell suspension samples (with SPE sample clean-up for the acid extractions).

#### 80:20 Methanol:Water (MeOH) Extraction

Cells were resuspended in 500 μL of 80:20 methanol:water (-80°C). The cell suspension was sonicated (5 x 0.5-second pulses at 50% intensity) to lyse cells, and the lysate was centrifuged at 17000 x g for 10 minutes at 4°C. The clarified extract was split between the PicoProbe^™^ and LC-MS methods (100 μL and 225 μL, respectively). For the LC-MS aliquot, 50 μL of acyl-CoA internal standard was added. The samples were dried under N_2_ and then resuspended in 50 μL of either Milli-Q water (PicoProbe™) or 5% SSA (LC-MS). 20 μL of the PicoProbe^™^ sample was added to duplicate wells of a white 96-well plate for use in the PicoProbe^™^ assay, which was performed per the manufacturer’s directions.

To evaluate the effect of the timing of internal standard addition during sample processing, cells were pelleted, resuspended in 1 mL of medium, and counted via a hemocytometer or a Coulter counter. To each of ten microfuge tubes, 200 μL of cell suspension was added, and the cells were pelleted and resuspended in 1 mL of 80:20 methanol:water (-80°C). 50-μL of acyl-CoA internal standard was added to half of the tubes (“early” samples), and all of the samples were sonicated and centrifuged as described above. To the tubes that had not received internal standard (“late” samples), 50 μl of acyl-CoA internal standard was added to the supernatant. All samples were dried under nitrogen, resuspended in 5% SSA, and analyzed via LC-HRMS as described.

#### Perchloric Acid (PCA) Extraction

Cells were resuspended in 500 μL of MS buffer (210 mM mannitol, 70 mM sucrose, 5 mM Tris-HCl, and 1 mM EDTA, pH 7.5). The cell suspension was deproteinized using the PCA-based deproteinization kit. Briefly, 100 μL of ice-cold PCA was added to the cell suspension, and the suspension was vortexed to mix. The cell suspension was sonicated (5 x 0.5-second pulses at 50% intensity) to lyse cells, and the lysate was centrifuged at 17000 x g for 10 minutes at 4°C. 30 μL of neutralization solution was added, and the suspension was mixed well and then placed on ice for 5 minutes. The suspension was centrifuged at 17000 x g for 1 minute at 4°C. The clarified extract was split between the PicoProbe™ and LC-MS methods (225 μL and 225 μL, respectively). 50 μL of the PicoProbe™ aliquot was added to duplicate wells of a white 96-well plate to be used directly in the PicoProbe™ assay, while 125 μL of the PicoProbe™ aliquot was applied to SPE columns. For the LC-MS aliquot, 50 μL of acyl-CoA internal standard was added prior to applying to SPE columns.

For the SPE sample cleanup, Oasis HLB SPE columns were used. The columns were preconditioned with 1 mL of methanol, followed by equilibration with 1 mL of Optima water. The sample aliquots were loaded onto the column and then washed with 1 mL of Optima water. Acetyl-CoA was eluted with 1 mL of 25 mM ammonium acetate in methanol. Samples were dried under N_2_ and resuspended in 50 μL of either Milli-Q water (PicoProbe™) or 5% SSA (LC-MS). 20 μL of the PicoProbe™ sample was added to duplicate wells of a white 96-well plate for use in the PicoProbe™ assay, which was performed per the manufacturer’s directions.

#### Trichloroacetic Acid (TCA) Extraction

Cells were resuspended in 500 μL of ice-cold 10% TCA. The cell suspension was sonicated (5 x 0.5-second pulses at 50% intensity) to lyse cells, and the lysate was centrifuged at 17000 x g for 10 minutes at 4°C. The suspension was centrifuged at 17000 x g for 1 minute at 4°C. The supernatant was split between the PicoProbe™ and LC-MS methods (225 μL and 225 μL, respectively). One hundred microliters of the PicoProbe™ aliquot was neutralized (∼ pH 8) using potassium hydroxide (2 M and 10 M) and 1 M Tris-HCl (pH 8) and then used directly in the PicoProbe™ assay (50 μL in each of two wells of a white 96-well plate), while 100 μL of the PicoProbe™ aliquot was applied to SPE columns. For the LC-MS aliquot, 50 μL of acyl-CoA internal standard was added prior to applying to SPE columns. The solid-phase extraction was performed as described for the PCA extraction.

To evaluate the effect of the timing of internal standard addition during sample processing, cells were pelleted, resuspended in 1 mL of medium, and counted via a hemocytometer or a Coulter counter. To each of ten microfuge tubes, 200 μL of cell suspension was added, and the cells were pelleted and resuspended in 1 mL of ice-cold 10% TCA. 50-μL of acyl-CoA internal standard was added to half of the tubes (“early” samples), and all of the samples were sonicated and centrifuged as described above. To the tubes that had not received internal standard (“late” samples), 50 μl of acyl-CoA internal standard was added to the supernatant. Solid-phase extraction and LC-MS analysis was performed as described.

### 96-Well Plate Metabolite Extraction and Sample Processing

We adapted our routine method for short-chain acyl-CoA extraction and sample processing for the use of 96-well plates and an automated pipetting system. The medium was aspirated from each well of a 96-well cell culture plate, and 100 μL of ice-cold 10% TCA was added to each well. A 100x dilution of the short-chain acyl-CoA internal standard was prepared, and 50 μL of the diluted internal standard was added to each well. The 96-well plate was mixed well by hand to prevent potential cross-contamination or loss of sample from vortexing or other vigorous mixing techniques. Samples were sonicated using 30 x 0.5 second pulses (50% intensity) via a sonicator equipped with an 8-tip probe. Protein and debris were precipitated by centrifugation at 2000 x g for 10 minutes at 4°C. The supernatant was transferred to a deep-well 96-well plate, which then was centrifuged for an additional 10 minutes at 2000 x g and 4°C to ensure that any precipitate that may have been transferred to the deep-well plate was forced to the bottom of the wells. Using a Tomtec Quadra4 liquid handling workstation, an Oasis HLB 96-well elution plate (30 mg of sorbent per well) was preconditioned and equilibrated with 1 mL of methanol and 1 mL of Optima water, respectively. Using the Tomtec Quadra4, the supernatant was applied to an Oasis HLB 96-well elution plate (30 mg of sorbent per well), the plate was washed with 1 mL of Optima water, and acetyl-CoA was eluted into a deep-well 96-well plate using 1 mL of 25 mM ammonium acetate in methanol. The eluent was evaporated to dryness under N_2_, and the samples were resuspended in 50 μL of 5% SSA using the Tomtec Quadra4. 15 μL of each sample was injected for LC-HRMS analysis.

### Minimum Cell Count Evaluation

The minimum cell count required to generate an adequate acetyl-CoA response via LC-HRMS was evaluated. HepG2, HAP1, or Panc-1 SLC25A20 cells were counted via a Coulter counter. For each cell type, cell suspensions were prepared at a concentration of 40000 cells per 100 μL in high-glucose DMEM with 10% FBS and 1% penicillin/streptomycin. Serial dilutions were prepared from the stock cell suspensions down to a concentration of 625 cells per 100 μL, after which 100 μL of each stock and serial-diluted cell suspension was added to each of four wells of a 96-well plate. The cells were incubated overnight, and metabolite extraction and sample processing were performed in a 96-well plate as described.

### Multiplexed Analysis of Short-Chain Acyl-CoAs

Short-chain acyl-CoAs were extracted from AML cell line and primary cell pellets. Cells were resuspended in 1 mL of ice-cold 10% TCA, and 50 μL of acyl-CoA internal standard was added. The cell suspension was sonicated (5 x 0.5-second pulses at 50% intensity) to lyse cells, and the lysate was centrifuged at 17000 x g for 10 minutes at 4°C. The suspension was centrifuged at 17000 x g for 1 minute at 4°C. SPE sample cleanup was performed as described. Serial dilutions of a short-chain acyl-CoA calibration standard mix (prepared in-house from commercially available acyl-CoAs) were prepared and processed in parallel with the samples. For the 11 short-chain acyl-CoAs where a standard curve was available, the amount (in pmol) of analyte in the sample was interpolated from the standard curve. The analyte amount was converted to cellular concentrations by accounting for cell volume and number of cells (Table S1). For each sample, the mean and standard deviation for each analyte were calculated via Microsoft Excel. For analytes lacking a standard curve (CoA-glutathione, hexanoyl-CoA, and malonyl-CoA), the relative abundance (peak area of analyte relative to the peak area of acetyl-CoA internal standard) was calculated and reported.

### LC-MS/MS Quantitation

For the LC-HRMS method, acetyl-CoA and other short-chain acyl-CoAs were analyzed as previously described (Frey *et al*., 2016) using an Ultimate 3000 quaternary ultra-high performance liquid chromatograph coupled to a Q Exactive Plus mass spectrometer (Thermo Scientific), with modifications for a two-column setup. A modified gradient was adopted using solvent A (5 mM ammonium acetate in water), solvent B (5mM ammonium acetate in 95:5 (v:v) acetonitrile: water) and solvent C (0.1% formic acid in 80:20 (v:v) acetonitrile: water). For the triple quadrupole LC-MS/MS method, modifications to accommodate binary solvent delivery and a triple quadrupole mass analyzer (Waters Acquity UPLC with a binary pump coupled to a Thermo Scientific TSQ Vantage mass spectrometer) were made. Data was acquired and processed using Xcalibur version 4.3 and TraceFinder™ 5.1 software, respectively (Thermo Fisher Scientific).

### Statistical Analysis

All statistical tests were performed with GraphPad Prism (v. 9.4.1). Bias and analytical method agreement were assessed using Bland-Altman plots and Deming regressions, using default parameters and the LC-HRMS method as the reference assay.

## Results

### Elabscience ELISA kit is not able to measure acetyl-CoA in some commercially available standards, cells, and tissue extracts

To test the ability of the Elabscience ELISA kit to measure acetyl-CoA, metabolite extracts from HepG2 cells or mouse tissue (heart or skeletal muscle) were analyzed via the Elabscience acetyl-CoA ELISA kit in parallel with LC-MS analyses. A standard curve was generated using the supplied acetyl-CoA material; however, an attempt to generate a standard curve using commercial acetyl-CoA (Sigma-Aldrich) that we use for preparing LC-MS standard solutions was unsuccessful (Fig. S1A). Using the ELISA kit, only four cell samples exhibited a response above that of the lowest-level standard (LLOQ, 0.04 pmol/well) (Fig. S1B). None of the tissue samples exhibited a response above that of the LLOQ (Fig. S1C). Based on the overall mean acetyl-CoA concentration determined for HepG2 cells via LC-HRMS (52 pmol/million cells) (Fig. S1D), the sample extracts from 0.1 million cells were expected to produce a response within the quantitative range of the ELISA. Similarly, based on the concentrations of acetyl-CoA determined via LC-HRMS for the mouse tissue samples (Fig. S1E), the mouse tissue extracts were expected to contain acetyl-CoA levels within or above the quantitative range of the ELISA. These data indicated that the ELISA kit is not usable for the quantitation of acetyl-CoA in metabolite extracts from HepG2 cell and mouse tissue samples.

### Fluorometric PicoProbe™ assay is variably inconsistent with LC-MS methods across cell and tissue matrices and extraction methods

To compare acetyl-CoA quantitation between the PicoProbe™ and LC-MS assays, metabolite extracts were prepared from HepG2 cell or mouse heart tissue samples and applied to the PicoProbe™ assay or LC-MS assays (Fig. 1A). Three different extraction methods were compared as outlined in the methods section: 80:20 methanol:water, 10% TCA in water, and PCA. A comparison of LC-HRMS and LC-MS/MS assays also was performed.

**Figure 1.**
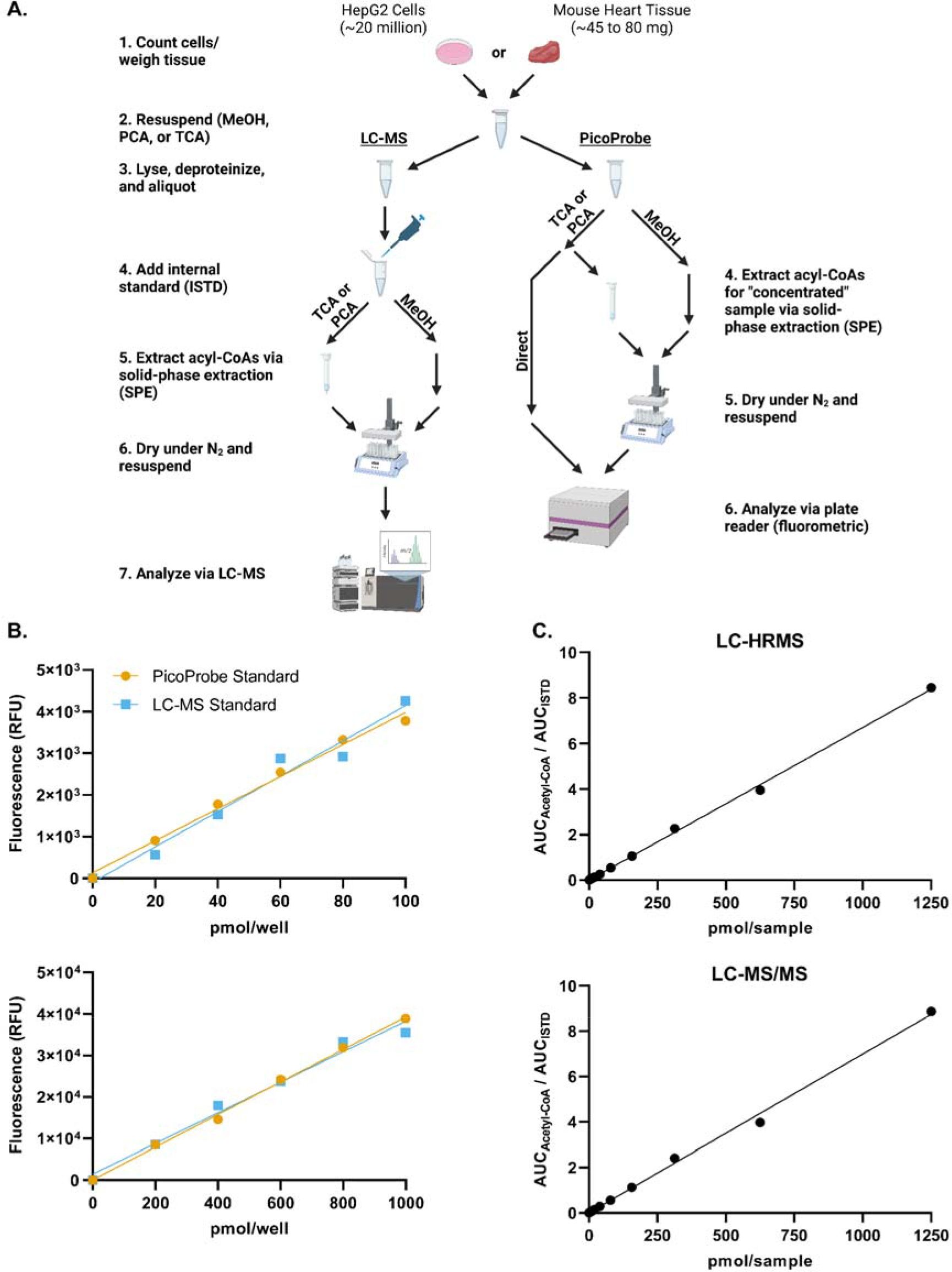
Method comparison schematic and calibration curves. (A) Outline of the method comparison experiment. (B) Standard curves were generated using the PicoProbe™ assay kit. The PicoProbe™ assay standards were prepared per the manufacturer’s recommendations. A set of standards also was prepared at the same concentrations using the stock standard used for the LC-MS assays. (C) Standard curves were generated via LC-HRMS and LC-MS/MS.

The PicoProbe™ kit supplies users with acetyl-CoA standard material for preparation of a standard curve, whereas we utilize commercial acetyl-CoA (Sigma-Aldrich) for LC-MS standard preparations. To ensure that no bias would be introduced into the method comparison study by using two different acetyl-CoA stock standard solutions (one for LC-MS and one for PicoProbe™), standard curves were generated via the PicoProbe™ kit using either the supplied acetyl-CoA standard or our LC-MS stock standard (Fig. 1B). Both standard materials produced similar standard curves, which was confirmed by Deming regression analysis (Fig. S2A-S2B). Standard curves generated via LC-HRMS and LC-MS/MS were similar (Fig. 1C and Fig. S2C). Thus, we concluded that the PicoProbe™ and Sigma-Aldrich acetyl-CoA standards are comparable and that the assays produce similar calibration curves from neat standards across the range used.

Acetyl-CoA was extracted from HepG2 cells or mouse heart tissue using three different extraction methods and analyzed via the PicoProbe™ assay, LC-HRMS, and LC-MS/MS (Fig. 2A). The PicoProbe™ manufacturer’s protocol recommends PCA deproteinization, followed by neutralization, and addition of the neutralized extract to the 96-well plate. Using this protocol (“PP – Direct” in Fig. 2A), the measured concentration of acetyl-CoA in HepG2 cells was 12 pmol/million cells, which was 43% of the concentration determined via LC-HRMS (28 pmol/million cells). For the LC-MS assays, the neutralized PCA extract was applied to a solid-phase extraction column for sample clean-up prior to LC-MS analysis. Therefore, we investigated if this same strategy could be utilized for the PicoProbe™ assay (“PP – Concentrated”). The acetyl-CoA concentration in the solid-phase extracted samples was 14 pmol/million cells, similar to the sample extract analyzed directly (12 pmol/million cells). The sample background fluorescence was lower for the samples passed through the SPE columns (Fig. S3C), potentially due to requiring a smaller volume of the more concentrated sample extract for analysis (20 μL compared to 50 μL). The PicoProbe™ assay did not generate usable data for the mouse heart tissue, perhaps due to high sample background fluorescence (Fig. S3E), which generally was observed across all cell and tissue samples (Fig. S3).

**Figure 2.**
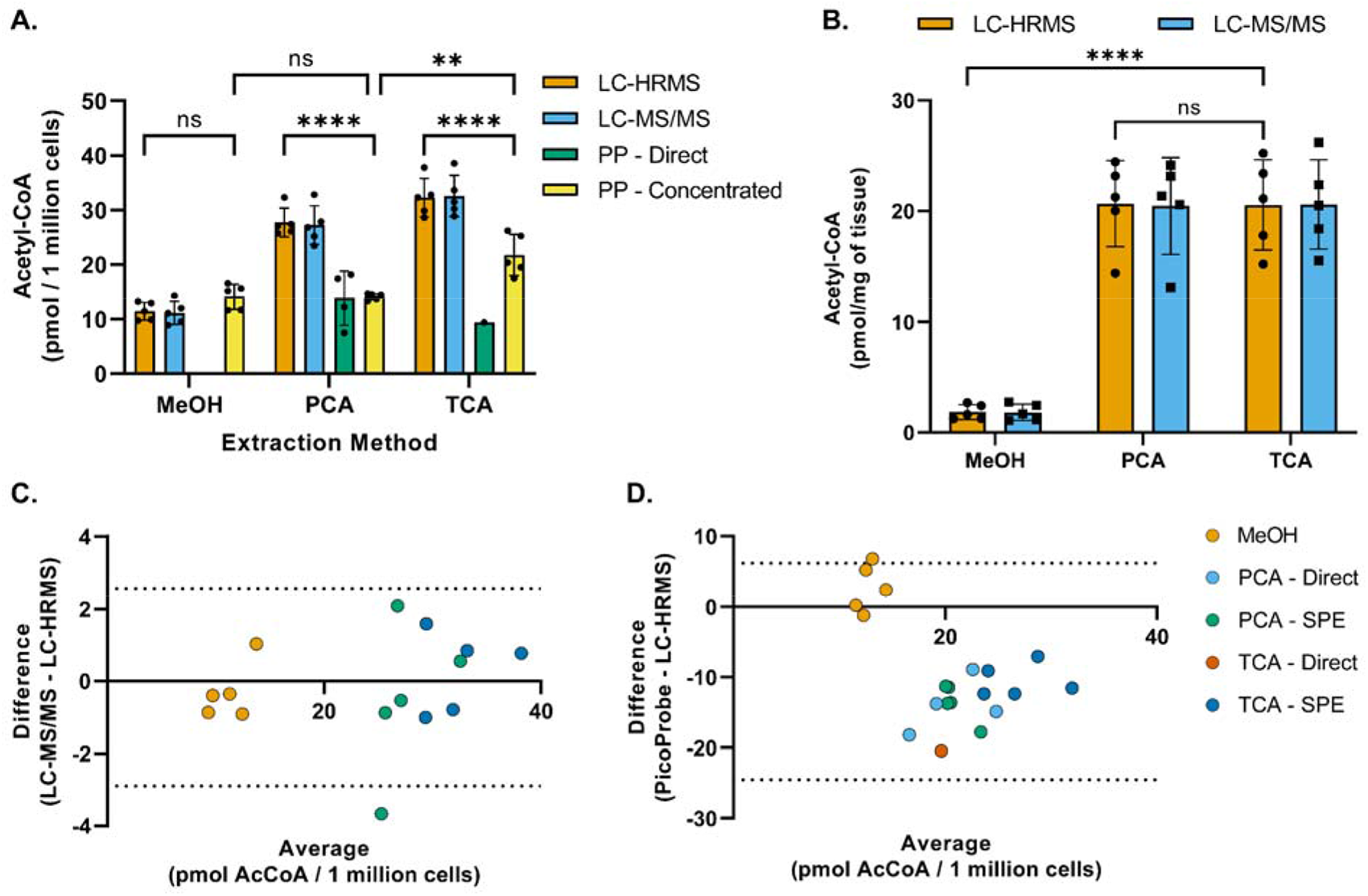
Method comparison of acetyl-CoA quantitation techniques and extraction methods. (A) Acetyl-CoA was extracted from HepG2 cells using -80°C 80:20 methanol:water (MeOH), perchloric acid (PCA) or 10% trichloroacetic acid in water (TCA) as described in the methods section and then analyzed via LC-HRMS, LC-MS/MS, or the PicoProbe™ (PP) assay. Each symbol represents an individual replicate sample, and error bars represent standard deviations. Statistical comparisons were performed via two-way ANOVA with post-hoc Tukey’s correction for multiple comparisons. The “PP – Direct” data were excluded from statistical comparisons due to the lack of replicates exhibiting a signal above the lower limit of quantitation. For each extraction method, no statistical significance was observed between the means of the LC-MS techniques. (B) Acetyl-CoA was extracted from heart tissue samples as described in the methods section and then analyzed via LC-HRMS or LC-MS/MS. For each extraction method, each symbol represents tissue from one of five different mice, and error bars represent standard deviations. Statistical comparisons were performed via two-way ANOVA with post-hoc Tukey’s correction for multiple comparisons. For each extraction method, no statistical significance was observed between the means of the LC-MS techniques. (C and D) Bland-Altman plots comparing sample results generated via (C) the LC-MS/MS assay or (D) the PicoProbe™ assay to the sample results generated via the LC-HRMS assay. **p≤0.01, ****p≤0.0001, ns = no significance

For the TCA extractions, only one of the five neutralized TCA extracts generated a background-corrected signal above that of the LLOQ when used directly for the PicoProbe™ assay (Fig. 2A). The acetyl-CoA concentration in the TCA extracts that were applied to SPE columns was 22 pmol/million cells, which was significantly lower than that determined via LC-HRMS (32 pmol/million cells) (Fig. 2A, 2D). Again, the SPE clean-up allowed for use of a smaller volume of sample extract and reduced the sample background fluorescence (Fig. S3D). When using methanol extractions, the acetyl-CoA concentration was 14 pmol/million cells, which was similar to that observed for the direct and solid-phase extracted PCA extracts (12 and 14 pmol/million cells, respectively) (Fig. 2A). The methanol extracts produced high sample background fluorescence relative to the other extraction methods (Fig. S3B).

Across all extraction methods, the LC-HRMS and LC-MS/MS assays produced similar acetyl-CoA measurements (Fig. 2A-2C). However, for both LC-MS methods, acetyl-CoA concentrations were significantly lower in cold methanol extracts than in acid extracts. For the purposes of the method comparison study, the LC-MS internal standard was already prepared in TCA and added after sonication during sample processing. In our typical LC-MS sample processing scheme, the internal standard is added prior to sonication to account for potential analyte loss during this step. To investigate if this modification to sample processing had a differential effect on the quantitation of acetyl-CoA in the cold methanol extracts, acetyl-CoA was extracted from HepG2 cells using either cold methanol or TCA, with the internal standard added either before (“early”) or after (“late”) the sonication step. Consistent with the initial results, cold methanol extraction produced lower acetyl-CoA concentrations than those observed using TCA extraction (Fig. S4). The timing of internal standard addition did not affect the TCA extracts, whereas there was a slight trend of lower acetyl-CoA concentrations with the “late” cold methanol extracts compared with the “early” cold methanol extracts. A possible explanation for this would be residual acetyl-CoA bound to protein in the cold methanol extraction that was liberated in the PCA and TCA extractions as has been reported for other assays [16-19]. We did not examine this further, but analysts should be aware of this trend for comparison between values reported by different extraction methods.

### Internal standardization improves the linear range of LC-MS assays

Since acetyl-CoA concentration changes in response to nutrient environment, cell signaling, and genetic background, the linear dynamic range of an assay may be an important consideration. We investigated the linear dynamic range of LC-HRMS and LC-MS/MS assays of acetyl-CoA with or without using a ^13^C_3_,^15^N_1_-acetyl-CoA internal standard. We found that internal standardization improved precision of calibration in both LC-HRMS and LC-MS/MS (Fig. S5A and B) and improved linearity across calibration and sample concentration ranges for both LC-HRMS and LC-MS/MS, with LC-HRMS benefitting more from normalization to an internal standard (Fig. S5C and D).

### Precise and linear acetyl-CoA/ISTD response ratio can be achieved at low cell numbers

Being able to detect and quantitate acetyl-CoA at low cell numbers would allow for higher throughput sample processing, as well as flexibility with regards to cell treatment protocols. Toward that end, we adapted our sample processing workflow to work with 96-well plates and the Tomtec Quadra4 liquid handling workstation. Utilizing this approach, we evaluated the minimum number of cells required to achieve a precise acetyl-CoA/ISTD ratio via LC-HRMS analysis for three different cell lines. For HepG2 cells, as few as 5000 cells generated a response ratio above background levels with acceptable precision (%CV ≤ 20), and a linear response ratio up to 40000 cells was observed (Figure 3A-3C). Similar results were obtained with Panc-1 SLC25A20 knockout cells, with precise response ratios above background achieved starting at 10000 cells (Fig. S6A-S6C). The response ratio of acetyl-CoA in HAP1 cells was more variable (%CV > 20% at all cell numbers) and exhibited less linearity across the full range of cell numbers evaluated (Fig. S6D-S6F). These results likely were due to the lower response ratio observed in these cells compared with the other cell lines tested. Together, these data indicate that quantitation of acetyl-CoA via LC-HRMS can be achieved when using low cell numbers and a workflow adapted for an automated pipetting system.

**Figure 3.**
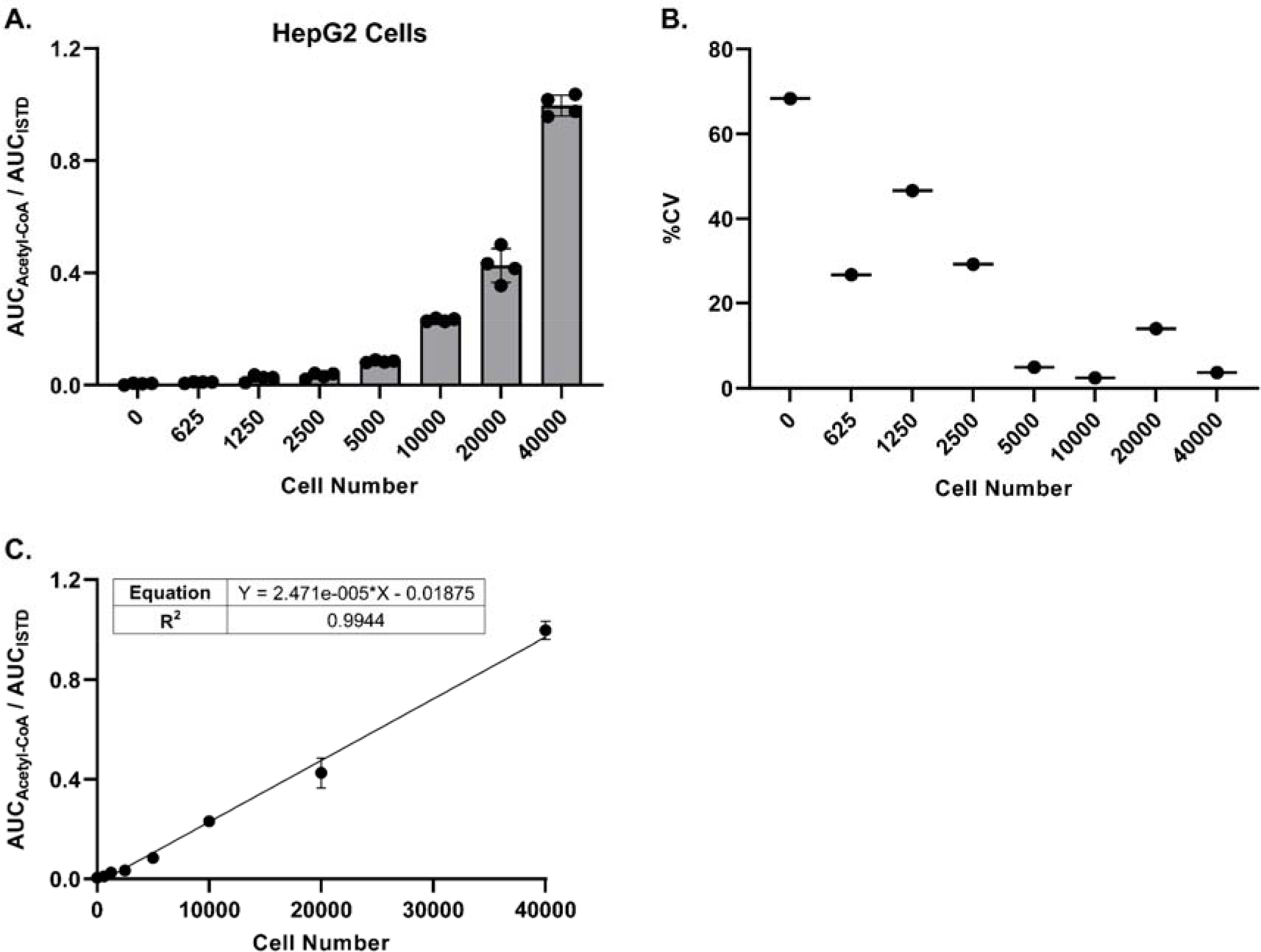
LC-HRMS analysis enables relative quantitation of acetyl-CoA at low cell numbers. HepG2 cells were plated in a 96-well plate at various cell numbers (n=4 wells per cell number), incubated overnight, and processed directly for LC-HRMS analysis. **(A)** Ratio of the peak area (AUC) of acetyl-CoA versus the peak area of the acetyl-CoA internal standard. Each symbol represents an individual replicate, and error bars represent standard deviations. **(B)** Percent coefficient of variation of the individual replicates (n=4) at each cell number. Some replicates at the lower cell numbers did not have a detectable acetyl-CoA response. **(C)** Linear regression of the peak area ratios. Each symbol represents the mean of individual replicates (n=4). Error bars represent standard deviations.

### LC-MS can be leveraged for multiplexed analysis of short-chain acyl-CoAs

An advantage of LC-MS assays over commercial kits is the ability to measure multiple analytes simultaneously within the same sample and analytical run (multiplexing). We tested the multiplexing capability of our LC-HRMS method by measuring a panel of short-chain acyl-CoAs in 27 AML cell lines and 9 primary AML cells (Table S1). Absolute quantification was achieved for 11 short-chain acyl-CoAs, and the relative abundance was calculated and reported for 3 additional short-chain acyl-CoAs for which a standard curve was unavailable. Acyl-CoA concentrations were calculated as indicated in the Methods section and are included in Table S1.

## Discussion

The primary limitation of LC-MS based methodologies is the cost of the instrumentation itself and the expertise to use the instrumentation. Unlike LC-MS methods, commercial assay kits detect acetyl-CoA via colorimetric or fluorometric methods that require equipment that is more widely available to most laboratories (i.e., a plate reader with appropriate filters). However, currently available commercial kits are limited to the single-analyte quantitation of acetyl-CoA. Despite the limited scope of commercial kits, the ease of use and non-specialized equipment required could provide researchers with a quick and effective alternative to LC-MS methods, provided the only analyte of interest is acetyl-CoA. However, no comparison between commercial assay kits and LC-MS assays has been documented. In this study, we compared the ability of two commercial assay kits (Elabscience ELISA and BioVision/abcam PicoProbe™) to measure acetyl-CoA to that of two different LC-MS assays.

The ELISA-based acetyl-CoA assay produced interpretable data for only a neat acetyl-CoA standard from the manufacturer. Thus, the comparisons we could conduct were very limited. Extensive pre-purification of a biological sample may provide interpretable values, but we did not test this as it was not indicated in the manufacturers protocol.

The PicoProbe™-based assay has been widely reported for measurement of acetyl-CoA. These reports do include measurements from tissue samples where we were unable to obtain a measurement above the anticipated LLOQ based on the background and calibration curve conducted per the manufacturer’s directions. Again, we anticipate that extensive pre-purification of the sample may be able to provide interpretable values as SPE of samples before the PicoProbe™ assay did lower the background readings. It may be useful for users of the PicoProbe™ assay to report LLOQ and background levels as this may be a tissue and experiment specific issue.

LC-MS/MS methods provide highly sensitive and specific quantitation for a variety of metabolites (including acetyl-CoA), especially when utilizing appropriate internal standards [14, 20, 21]. In addition, LC-MS/MS methods used for acetyl-CoA quantitation typically are compatible with the simultaneous detection and quantitation of a variety of acyl-CoAs, allowing multiplexing with LC-MS based methods [3, 22]. In this study, we demonstrated multiplexing by measuring a panel of short-chain acyl-CoAs in a variety of AML cells. Another benefit of mass spectrometry-based assays is the ability to incorporate isotope analysis. In this project, we utilized a stable isotope-based internal standard incorporating 13C_3_,15N_1_-pantothenate into the coenzyme A portion of acetyl-CoA. This improved the linearity of calibration curves, especially of the LC-HRMS-based assay. Since none of the commercially available kits can incorporate isotopic information, we could not compare the benefits of isotope dilution in the kits.

This work focused on acetyl-CoA, but commerical kits for CoA and malonyl-CoA have also been reported. At the time of writing not all of these kits were available, as the malonyl-CoA kit had been withdrawn. Similarily, HPLC-UV [23-28] and HPLC-flourometric [29, 30] assays of underivatized and derivatized acyl-CoAs have been reported. Future work comparing these assays may be useful to the field in understanding the most accessible and appropriate assay for a given biological question.

## Acknowledgements

This work was supported by NIH grants T32GM142606 (DSK), R01DK109100 (ES), R01CA259111 and R01GM132261 (NWS), R01CA254354 and R21CA256128 (MWD).

## Figures (in order of appearance)

**Supplemental Figure 1.**
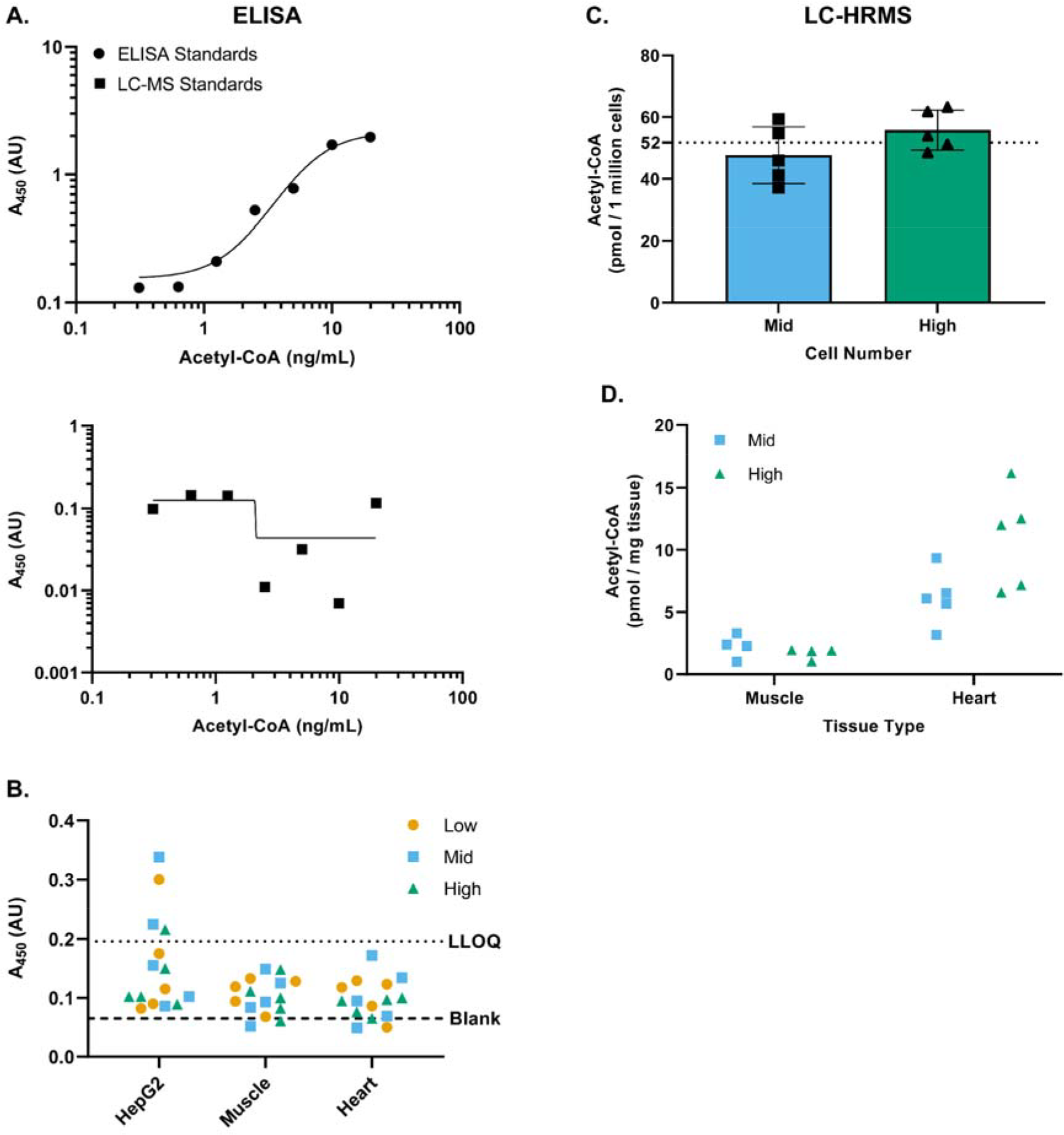
Acetyl-CoA ELISA kit does not provide reliable quantitation of acetyl-CoA in cell or tissue samples. Acetyl-CoA was measured in HepG2 cells and tissue samples (mouse skeletal muscle and heart) via an Elabscience acetyl-CoA ELISA kit or via LC-HRMS. For cell samples, 100 μL of 1-mL metabolite extracts from 10 million (high), 1 million (mid), or 100,000 (low) cells was analyzed. For tissue samples, 100 μL of 1-mL extracts of ∼30 mg (high), ∼10 mg (mid), or a 1:10 dilution of the 10 mg extract (low) was analyzed. (A) Four-parameter logistic curves generated via ELISA using standards prepared from Elabscience ELISA (top) or LC-MS (Sigma-Aldrich) (bottom) acetyl-CoA stock standard solutions. Absorbance from the blank was subtracted from the absorbance of each standard prior to plotting per the protocol. (B) Raw absorbance values of cell and tissue samples analyzed via ELISA. The lower limit of quantitation (LLOQ) (dotted line) refers to the raw absorbance of the lowest concentration standard (0.04 pmol/well), and the Blank (dashed line) refers to the raw absorbance of the blank, which did not contain added acetyl-CoA. Note that only a few cell samples exhibited a raw absorbance above that of the LLOQ absorbance, and the response did not scale with the number of cells utilized. (C) Acetyl-CoA measured in HepG2 cells via LC-HRMS. Each symbol represents an individual replicate sample, and error bars represent standard deviations. The overall mean (n=10) of the mid and high samples was 52 pmol/million cells. The low samples exhibited a response below the lower limit of quantitation. (D) Acetyl-CoA measured in mouse skeletal muscle and heart tissue via LC-HRMS. Each symbol represents tissue from one of four (skeletal muscle) or five (heart) different mice. The majority of the low samples exhibited a response below the lower limit of quantitation and, therefore, are not shown.

**Supplemental Figure 2.**
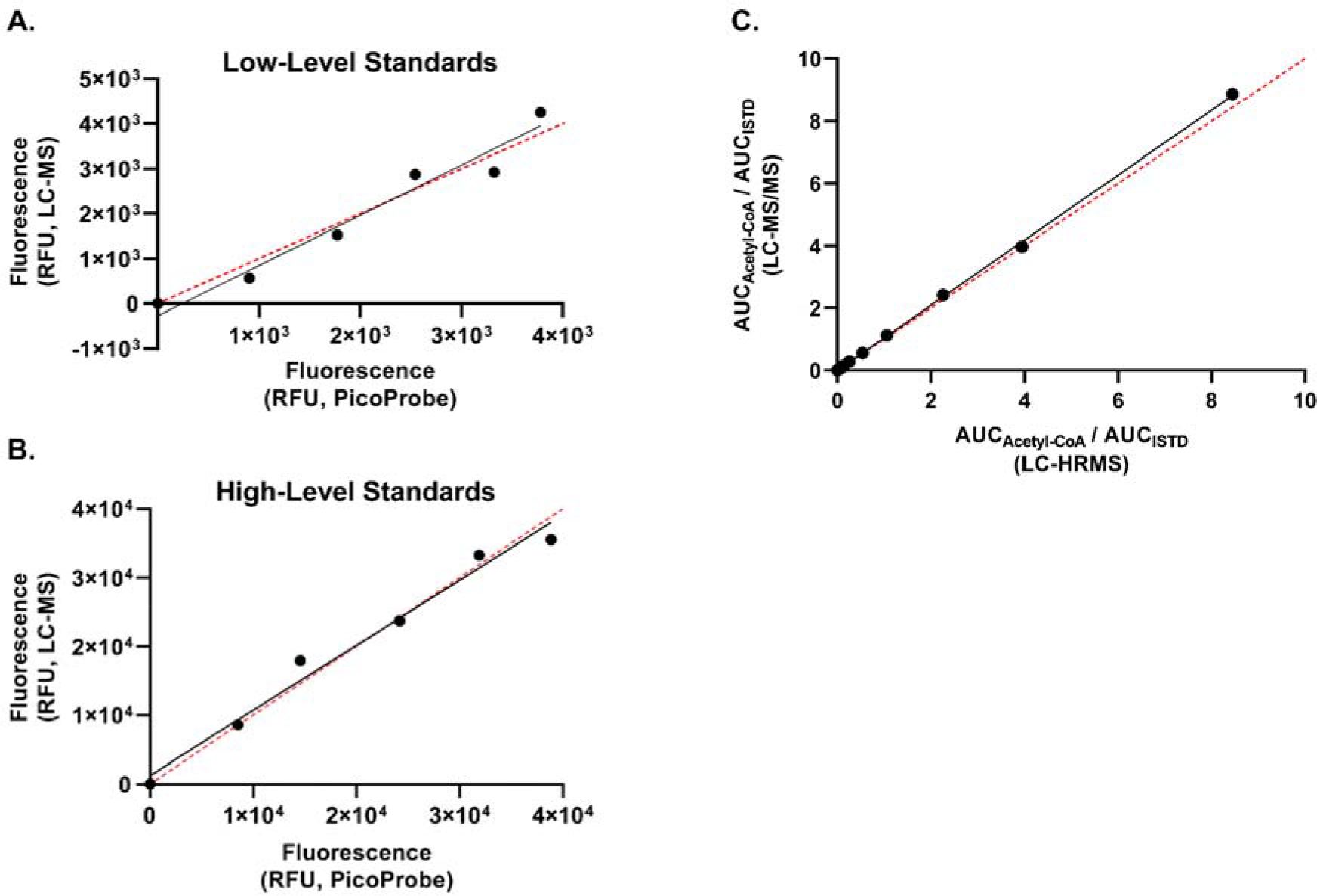
Deming regression analysis of PicoProbe™ and LC-MS standards analyzed via the fluorometric PicoProbe™ kit. Deming regression analysis was performed for (A) low- and (B) high-level standard curves generated using the PicoProbe™ assay kit. The PicoProbe™ assay standards were prepared per the manufacturer’s recommendations. A set of standards also was prepared at the same concentrations using the stock standard used for the LC-MS assays. The dashed line represents the y=x line of agreement. RFU = relative fluorescence units (C) Deming regression analysis was performed for standard curves generated via LC-HRMS and LC-MS/MS assays. The dashed line represents the y=x line of agreement. AUC = area under the curve

**Supplemental Figure 3.**
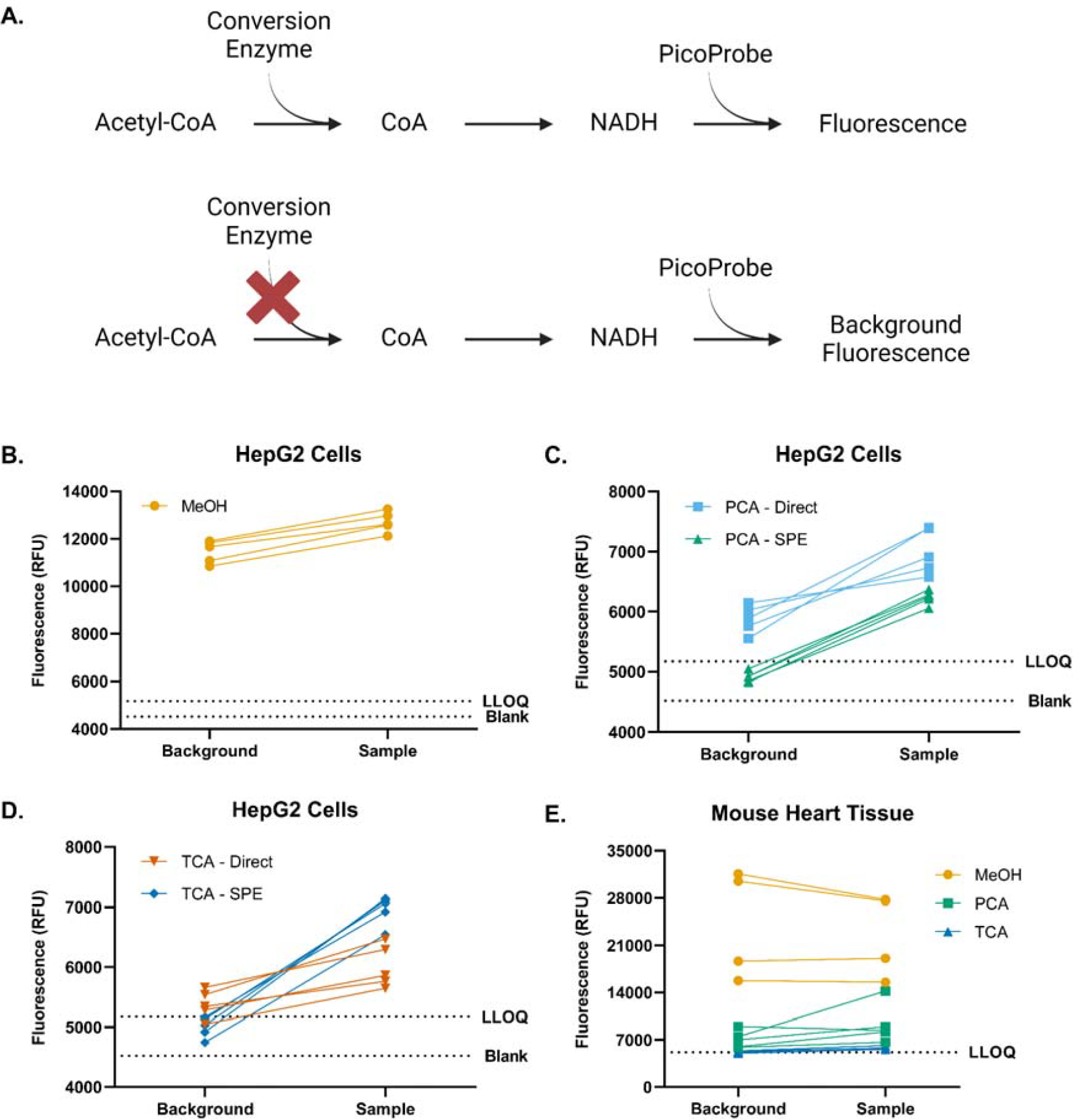
High background fluorescence interferes with PicoProbe™ measurements in mouse heart tissue samples. (A) Schematic of how background sample fluorescence is determined via the PicoProbe™ assay. The sample is added in equal volumes to each of two wells of a 96-well plate. To one well, a reaction mix containing the conversion enzyme is added. To the other well, a reaction mix omitting the conversion enzyme is added to evaluate background sample fluorescence. (B-D) Acetyl-CoA was extracted from HepG2 cells using (B) -80°C 80:20 methanol:water (MeOH), (C) perchloric acid (PCA) or (D) 10% trichloroacetic acid in water (TCA) as described in the methods section and then analyzed via the PicoProbe™ assay. Raw fluorescence from the background correction samples and the test samples was plotted, with lines connecting the corresponding background and test samples. (E) Acetyl-CoA was extracted from mouse heart tissue using MeOH, PCA, or TCA as described in the methods section and then analyzed via the PicoProbe™ assay. Raw fluorescence from the background correction samples and the test samples was plotted as in (B-D). Each symbol represents an individual replicate sample.

**Supplemental Figure 4.**
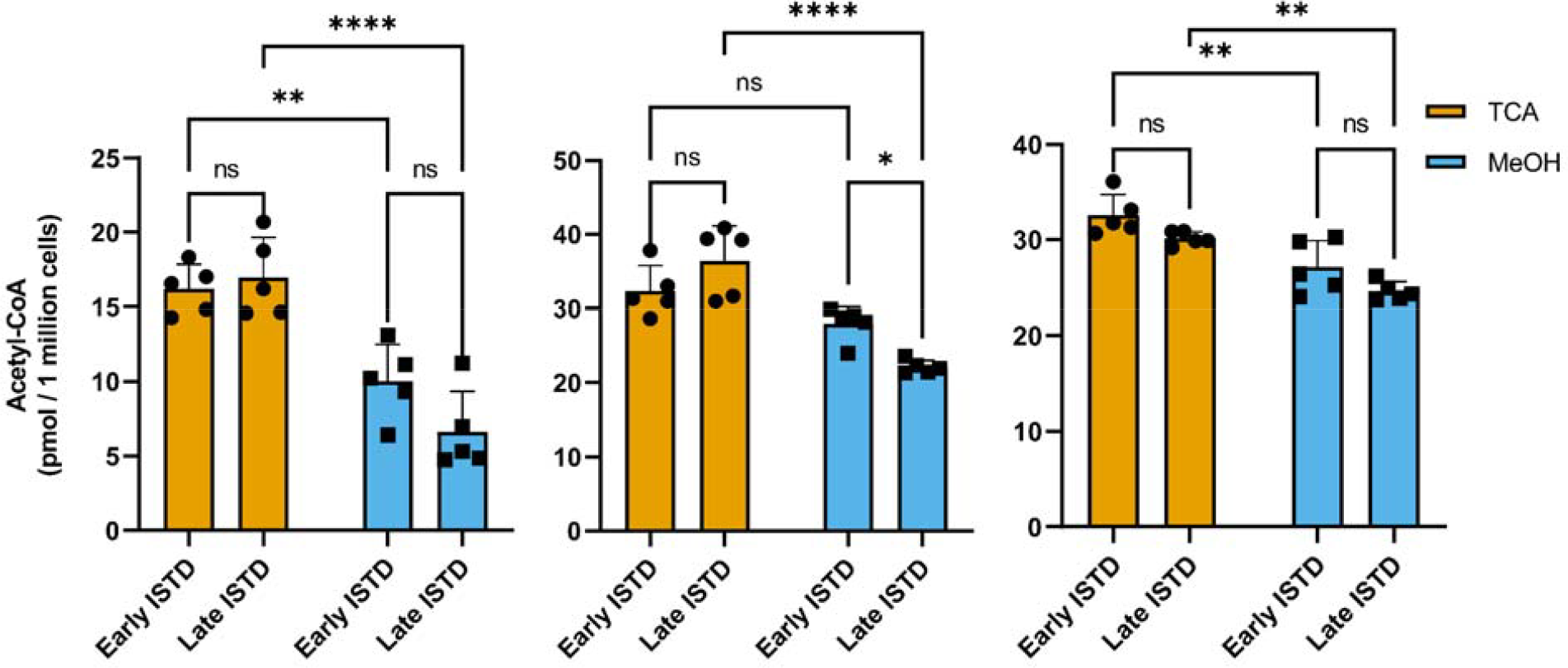
Acetyl-CoA concentration measured via LC-HRMS is affected by extraction method. Acetyl-CoA was extracted from HepG2 cells using -80°C 80:20 methanol:water (MeOH) or 10% trichloroacetic acid in water (TCA) as described in the methods section and analyzed via LC-HRMS. During sample processing, the internal standard was added either prior to sonication (“early ISTD”) or after sonication (“late ISTD”). Each symbol represents an individual replicate sample, and error bars represent standard deviations. Each graph represents an individual experiment. Statistical comparisons were performed via two-way ANOVA with post-hoc Tukey’s correction for multiple comparisons. *p≤0.05, **p≤0.01, ****p≤0.0001, ns = no significance

**Supplemental Figure 5.**
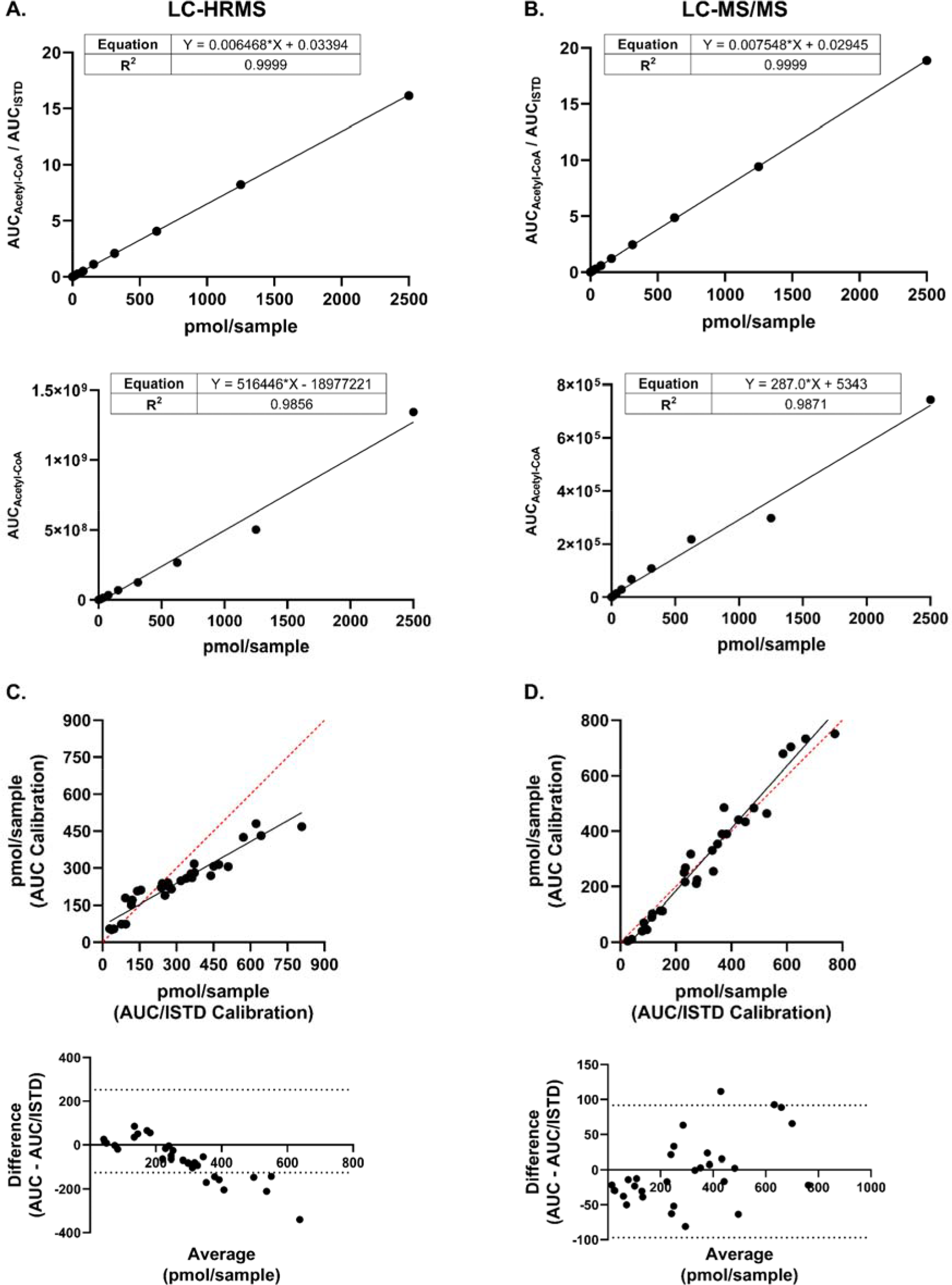
Using an internal standard for LC-MS analysis improves linearity. (A and B) Callibration curves were generated via (A) LC-HRMS or (B) LC-MS/MS analysis using either the ratio of the acetyl-CoA peak area (AUC) and the internal standard peak area (top) or only the acetyl-CoA peak area (bottom). (C and D) Deming regressions (top) and Bland-Altman plots (bottom) were generated from the sample results interpolated from the two different standard curves measured via (A) LC-HRMS or (B) LC-MS/MS analysis. The dashed line represents the y=x line of agreement. Data is representative of n=2 independent experiments.

**Supplemental Figure 6.**
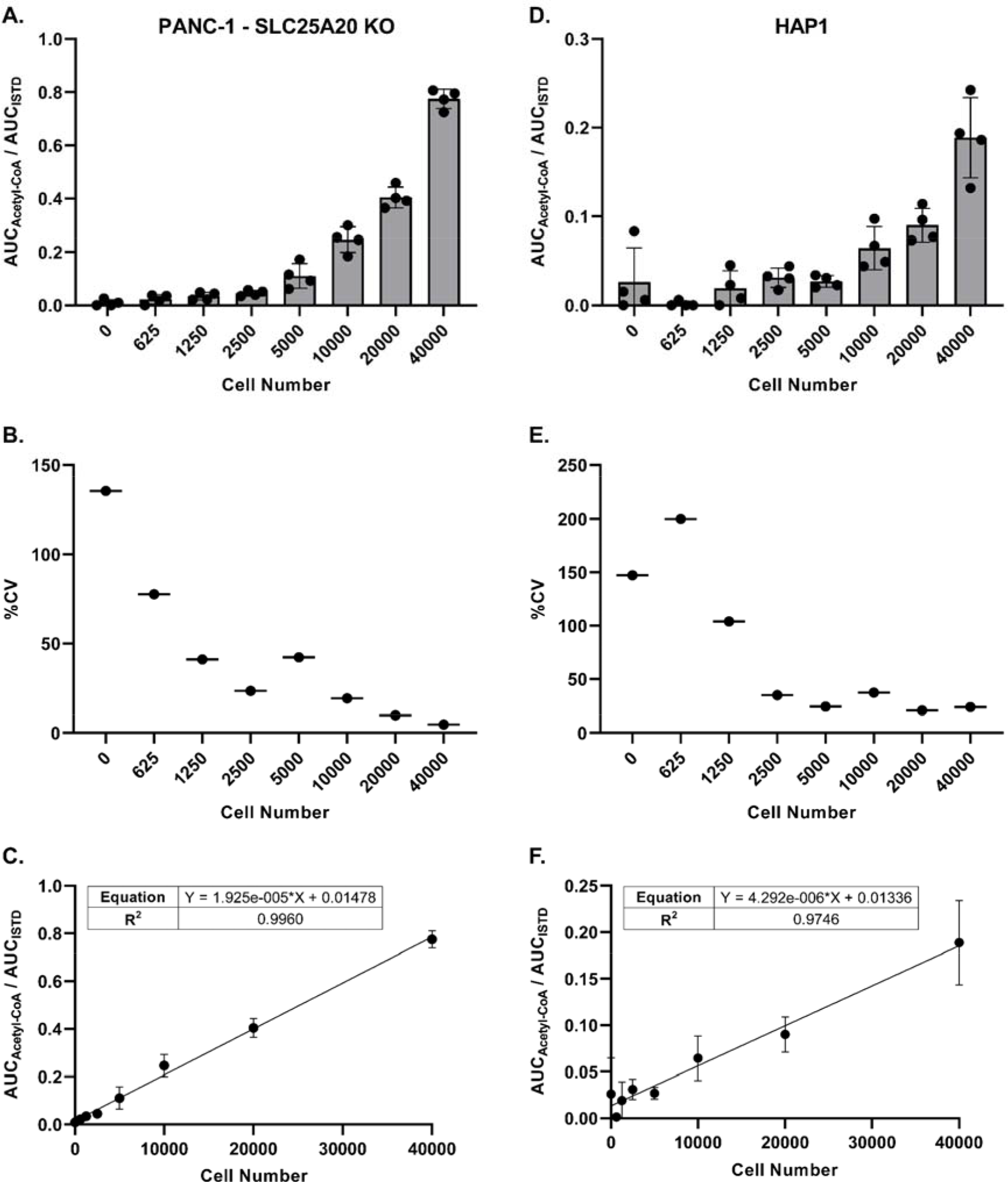
The ability of LC-HRMS analysis to achieve precise relative quantitation of acetyl-CoA at low cell numbers is cell-line dependent. Panc-1 SLC25A20 or HAP1 cells were plated in a 96-well plate at various cell numbers (n=4 wells per cell number), incubated overnight, and processed directly for LC-HRMS analysis. **(A and D)** Ratio of the peak area (AUC) of acetyl-CoA versus the peak area of the acetyl-CoA internal standard. Each symbol represents an individual replicate, and error bars represent standard deviations. **(B and E)** Percent coefficient of variation of the individual replicates (n=4) at each cell number. Some replicates at the lower cell numbers did not have a detectable acetyl-CoA response. **(C and F)** Linear regression of the peak area ratios. Each symbol represents the mean of individual replicates (n=4). Error bars represent standard deviations.

